# Constitutive activity of the inhibitory G protein pathway mediated by non-visual opsin Opn7b reduces cFos activity in stress and fear circuits and modulates avoidance behavior

**DOI:** 10.1101/2025.05.05.652173

**Authors:** Hanna Böke, Hannah Schulte, Maria Worm, Julia Bihorac, Brix Mücher, Martin Hadamitzky, Ida Siveke, Stefan Herlitze, Katharina Spoida

## Abstract

Constitutive activity of G protein-coupled receptors (GPCRs) plays an important role in brain function and disease including neurodegenerative and psychiatric disorders. The non-visual opsin Opn7b is a constitutively active G_i/o_ coupled GPCR which has been used to synchronize neuronal networks. Here we show that expression of Opn7b in the bed nucleus of the stria terminalis and the ventral tegmental area, two interconnected brain areas involved in modulating fear and stress responses, reduces the number of cFos positive neurons and modulates avoidance behavior in mice. Thus, by constitutively activating the Gi/o pathway Opn7b can be used as a tool to reduce cFos expression and to link cFos-expressing neurons to network- and pathway-specific behavior.

## Introduction

Constitutive activity of G protein-coupled receptors (GPCRs) refers to their ability to activate signaling pathways in the absence of an agonist. This phenomenon has important implications for receptor function, drug discovery, and the understanding of disease mechanisms. Constitutive activity has been observed in many GPCRs across different families and species and plays important physiological roles, such as providing tonic support for basal neuronal activity (Seifert and Wenzel-Seifert, 2002).

The midbrain dopamine system, which is crucial for processing rewards and associated stimuli such as anxiety, fear and stress, is tightly regulated by various GPCRs. Importantly, these receptors exhibit agonist-independent intrinsic constitutive activity that contributes to their regulation of the ventral tegmental area (VTA) and its projection targets including the bed nucleus of the stria terminalis (BNST) (Meye et al., 2014). By integrating information from various limbic structures, the BNST is a key structure in regulating and shaping motivated and anxiety related behavior (van de Poll et al., 2023). The BNST and VTA have a direct reciprocal connection, with the BNST serving as a major source of neuropeptide and GABAergic inputs to the VTA (Soden et al., 2022).

Among other brain areas various regions and cell types of the BNST and VTA are activated during stress and fear. Activation of neuronal circuits has been correlated with the induction of immediate early gene expression such as cFos (Chung, 2015; Chowdhury and Caroni, 2018). Acute stressors, such as restraint stress, increase cFos expression and neuronal activity in corticotropin-releasing factor (CRF) neurons in the dorsal BNST. This activation is consistent with the role of BNST CRF neurons in driving negative affective behaviors associated with stress (Maita et al., 2022). In addition, various forms of acute and chronic stress increases cFos expression and neuronal activity in different VTA neuron types including GABAergic and dopaminergic neurons (Lowes et al., 2021; Koutlas et al., 2022; Qi et al., 2022; Yu et al., 2022; Yang et al., 2023). This stress-induced activation of VTA neurons appears to play a role in stress-related behavior and may contribute to the development of anxiety-like behaviors (Qi et al., 2022; Mitten et al., 2024). Thus, elevated cFos expression by stress is correlated with increased network activity in the BNST and VTA. Therefore, decreased neuronal activity and following reduced cFos expression within these areas should modulate stress responses.

We recently characterized a non-visual opsin Opn7b from zebrafish in heterologous expression systems and mouse brain. Opn7b is a reverse photoreceptor, that constitutively activates the G_i/o_ pathway in the dark. This constitutive activity of Opn7b can be switched off by blue-green light (Karapinar et al., 2021). Opn7b is not expressed in mammals but in the brain, eye and testis of adult zebrafish (Davies et al., 2015). Activation of G_i/o_ coupled GPCRs such as inhibitory DREADDs (designer receptors exclusively activated by designer drugs) reduce cFos expression (Armbruster et al., 2007; Bruzsik et al., 2021; Tiwari et al., 2022). Thus we tested whether Opn7b without supplying exogenous ligands or drugs, could reduce cFos expression in anxiety- and fear-related circuits, and thereby affect anxiety- and fear-related behavior. Here we demonstrate that viral-mediated expression of Opn7b in the BNST and the VTA reduces cFos expression and modulates of odor-induced avoidance and fear responses.

## Materials and methods

### Subjects

Adult (>8 weeks old) male C57Bl/6 mice (stock No. #000664, Jackson Laboratory) were used in the present study. Mice were group housed, if possible (2-3 mice/cage), with a 12h light/dark cycle and constant room temperature. Food and water were available *ad libitum*. The procedures were performed during the light phase. All experiments were approved by the local ethics committee (Bezirksamt Arnsberg) and animal care committee of Nordrhein-Westfalen (LANUV; Landesamt für Umweltschutz, Naturschutz und Verbraucherschutz Nordrhein-Westfalen, Germany; AZ: 81-02.04.2021.A412). Studies were conducted in accordance with the European Communities Council Directive of 2010 (2010/63/EU) for care of laboratory animals and supervised by the animal welfare commission of the Ruhr-University Bochum.

### Stereotaxic surgery

2/1 pseudotyped adeno-associated viruses (AAV) carrying either the opsin parapinopsin or the constitutively active, G_i/o_-coupled optogenetic tool Opn7b were administered by stereotaxic application. Both optogenetic tools are fused to enhanced green fluorescent protein and expressed under control of the CMV promotor. Mice were initially anesthetized using 5 % isoflurane (v/v) whereas isoflurane levels were adjusted to 1.5-2.5 % (v/v) for the entire surgery. Carprofen (2 mg/kg) and buprenorphine (0.1 mg/kg) were injected subcutaneously for analgesia. For local anesthetic, lidocaine was applied to the scalp. Virus injection was conducted via pressure injection with customized glass pipets (tip diameter 5-10 µm). For injection into the BNSTad and VTA the following stereotactic coordinates were used, related to Bregma: BNSTad AP +0.26mm, ML +/-0.75mm, DV -4.2 mm and VTA AP - 2.9mm, ML +/-0.3-0.5 mm, DV -4.8mm. The viral construct were expressed for approximately two weeks prior to behavioral testing, as this time frame was shown to be sufficient to induce maximal effect on network modulation (Karapinar et al., 2021).

### Behavioral testing

The experimental paradigm used was based on a behavioral paradigm established by (Bruzsik et al., 2021). Mice were tested in a behavioral paradigm to investigate innate fear responses using a synthetic analog to a fox anogenital product (2-methyl-2-thiazoline; 2MT). Prior to the behavioral experiments, the mice were handled for 2-3 days. Behavioral testing was carried out in an open field arena (50 × 50 × 38 cm) with a light intensity of ∼1500 lux in the middle. The 2MT (cat. No. L01329.14, Thermo Fisher (Kandel)) was diluted 1:50 and presented on a filter paper placed in a glass petri dish in one corner. The area around the odor source was defined as the “odor zone” (12.5 × 12.5 cm). Testing was repeated on two consecutive days with different 2MT conditions which were characterized by the amount of 2MT used (according to (Bruzsik et al., 2021); low 2MT: 10 µl, high 2MT: 250 µl). At the start of the behavioral test the mice were placed in the corner opposing the odor zone and were left to freely roam and explore the arena for 10 minutes. Between individuals, the filter paper was removed, and the arena was cleaned with water and 70 % EtOH and left to ventilate for 10 minutes. Four behavioral variables were used to characterize innate fear responses: (1) the locomotor exploratory activity, measured by the total distance moved in cm, (2) maximum velocity, (3) time spent in the odor zone, based on the center point of the animal, and (4) time spent freezing. All behavioral variables were quantified using Ethovision XT 15 software.

### Histology and immunohistochemistry

On the second day of behavioral testing, mice were deeply anesthetized and transcardially perfused with ice-cold PBS (1x) followed by 4 % paraformaldehyde in PBS (PFA, w/v, pH 7.4, Sigma Aldrich) 90 minutes after the test. Brains were postfixed in 4 % PFA for two hours and cryoprotected in 30 % sucrose (w/v, Fisher Scientific). For subsequent immunohistochemical stainings, brains were cut into 30 µm sections using a cryostat (CM3050 S, Leica). All stainings were performed free floating using a 24-well plate filled with TBS (50 mM Tris-HCl, 150 mM NaCl, pH 7.5). Sections were blocked with 10 % normal donkey serum (NDS, Merck Milipore) in 0.3 % TBS-Triton X-100 (TBS-T, triton X-100 in TBS) to reduce non-specific antibody binding. BNST sections were incubated with antibody for cFos (rabbit anti cFos, Cat. No. 226 008, Synaptic Systems) to label neuronal activity. VTA sections were incubated with antibody for cFos, and TH (mouse anti TH, MAB318, Merck Millipore) as a marker for dopaminergic neurons in the VTA. Primary antibodies were incubated in a solution of 3 % NDS in 0.3 % TBS-T, as well as the secondary antibodies. Secondary antibody solution included donkey anti rabbit Cy5 (Code no. 711-175-152, Jackson Immuno Research) for cFos and donkey anti mouse Dylight 550 (SA5-10167, Thermo Fisher Scientific) for TH.

### Imaging and Analysis

All immunolabeled brain sections were captured using a confocal laser scanning microscope (TCS SP5II, Leica Microsystems) by using 10x/0.3 NA and 20x/0.7 NA objectives. Z-stack images with 10 optical planes were taken. For quantitative analysis, images were processed using ImageJ software (Schneider et al., 2012). The quantification of c-Fos expression was confined to predefined regions of interest (ROIs), which were delineated using the ImageJ “Polygon Selection” tool. Within the BNST, cell counting was restricted to the BNSTad. A minimum of eight corresponding ROIs within the BNST and VTA from both hemispheres were analyzed. BNSTad ROIs were delineated using the lateral ventricle and the anterior commissure, while VTA ROIs were defined using TH as a marker for dopaminergic neurons. cFos immunoreactive cells were counted using a semi-automated quantification with the “Thresholding” and “Analyze Particles” functions (Bihorac et al., 2024). c-Fos-positive cells were defined based on a particle size range of 15–300 pixel^2^ and a circularity between 0.1 and 1.0. Autofluorescent artifacts, which were mistakenly counted as c-Fos immunoreactive cells, were manually excluded from the analysis.

### Statistics

Graphs were generated using SigmaPlot 12.5 (Systat Software) and the data were analyzed with IBM SPSS Statistics (Version 29.0) software. Normality (Shapiro-Wilk test) and homogeneity of variance (Levene’s test) were tested before each analysis. An unpaired Student’s t-test was utilized to calculate statistical significance. As a non-parametric alternative, the Mann-Whitney U test was employed. Statistical significance was determined using a critical alpha level of 0.05 (p ≤ .05).

## Results

To investigate whether the constitutive activation of the G_i/o_ pathway reduces cFos expression in neuronal circuits and if this reduction can be correlated with a change in behavior we expressed Opn7b, a constitutively active and as a control parapinopsin, a non-constitutively active GPCR, in neuronal circuits associated with stress-induced behaviors (Eickelbeck et al., 2019; Karapinar et al., 2021). We used a predator odor-evoked innate fear paradigm to test for threat anticipation and fear modulation under ambiguous threats (Bruzsik et al., 2021). 2-Methyl-2-thiazoline (2MT) is an innate fear inducer. It is a synthetic derivative of compounds found in fox anogenital secretions, which naturally evoke fear responses in prey animals (Apfelbach et al., 2015; Rosen et al., 2015).

We first injected male mice with AAV expressing Opn7b or parapinopsin under the CMV promotor in the BNST. Two weeks post-injection, we conducted behavioral tests comparing control mice, expressing parapinopsin, to those expressing Opn7b (Fig. 1A). Animals were exposed to water or to two different dosages of 2MT and various behavioral parameters including total distance moved, mean velocity of movement, time spend in odor zone, and freezing were analyzed (Fig. 1B-F). We found that Opn7b expressing mice in comparison to parapinopsin expressing mice revealed a decrease in the total distant moved at high 2MT concentrations (Fig. 1C) and a decrease in the time spend in the odor zone at low 2MT concentration (Fig. 1E). The decrease in time spend in the odor zone suggests an increase in avoidance behavior when Opn7b is expressed in the BNST (Albrechet-Souza and Gilpin, 2019; Weera et al., 2021). 90 min after the behavioral experiment mice were sacrificed and cFos expression was compared between parapinopsin and Opn7b expressing mice. We found that the number of cFos expressing cells in the BNST was reduced by 33% in Opn7b expressing mice in comparison to parapinopsin expressing mice (Fig. 1. G,I) with no change in cFos expression in the VTA (Fig. 1 H,J).

**Figure 1:**
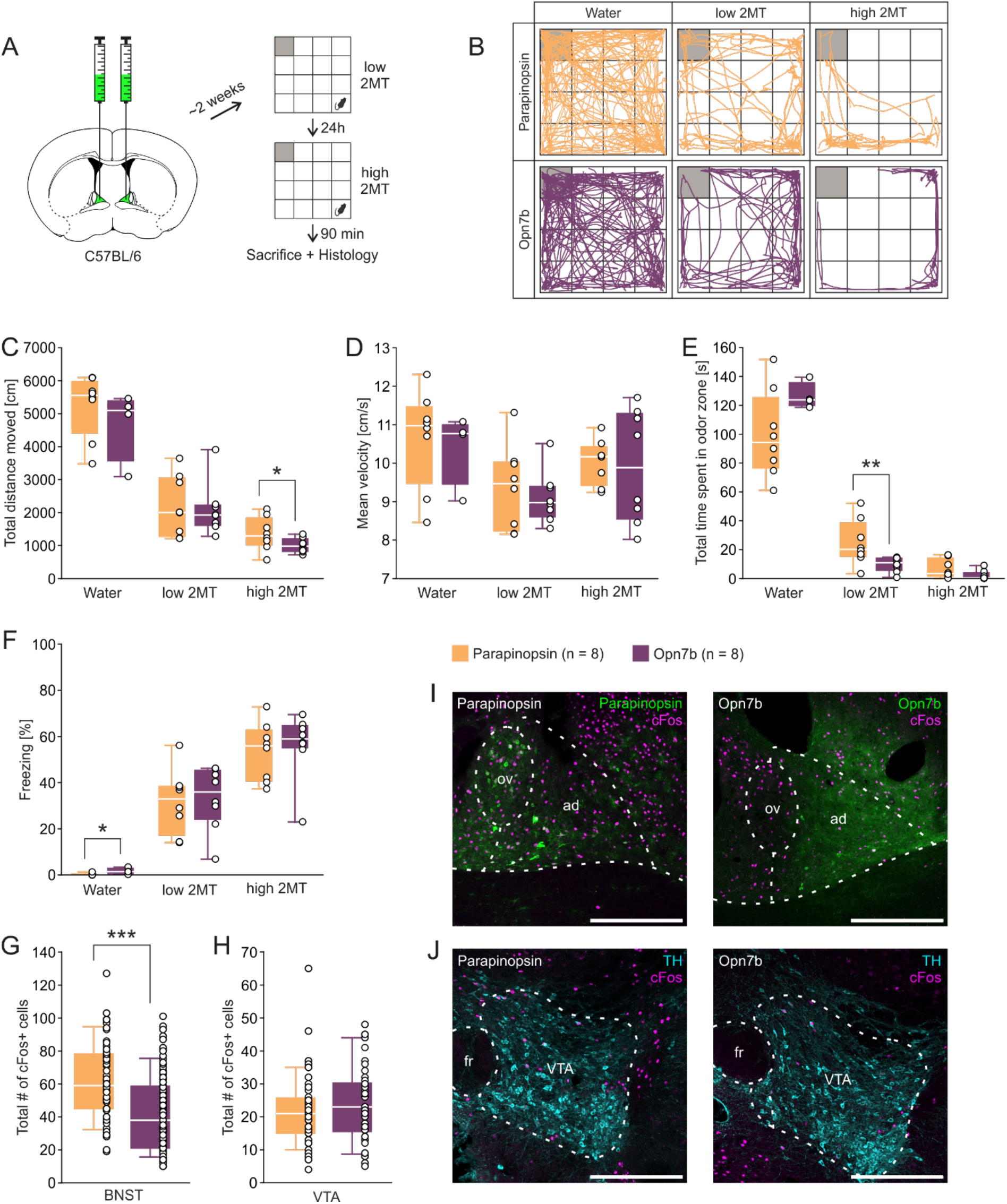
Injection of the constitutively active optogenetic tool Opn7b into the BNST reduces cFos expression in the BNSTad. **A)** Stereotactic injection of the viral constructs encoding either Opn7b or parapinopsin into the anterodorsal nucleus of the bed nucleus of the stria terminalis (BNSTad) of C57BL/6 mice followed by behavioral testing in the 2-methyl-thiazoline (2MT) paradigm with the grey square representing the odor zone. **B)** Exemplary trajectory plots of individual mice of both groups exposed to Water as well as low and high 2MT condition in an open field arena. **C)** Mice expressing Opn7b in the BNST show a decreased total distance moved compared to mice expressing parapinopsin in the high 2MT condition (t_(14)_ = 1.84, p = .044). **D)** No differences between groups in the mean velocity in all conditions were found. **E)** Opn7b injection decreased the time spent in the odor zone only during low 2MT exposure (U = 7.00; z = -2.626, p = .007, r = 0.66) but not high 2MT exposure or the Water condition. **F)** Mice expressing Opn7b in the BNST showed less freezing in the Water condition (U = 28.00; z = 2.075, p = .048, r = 0.60) compared to mice expressing parapinopsin. **G)** The mice expressing Opn7b (quantified areas: n = 96) show a decrease in the total number of cFos immunoreactive cells in the BNSTad (U = 1667.50; z = -5.116, p < .001, r = 0.40) compared to mice expressing parapinopsin (quantified areas: n = 66). **H)** There were no differences in the total number of cFos immunioreactive cells in the ventral tegmental area (VTA) between the groups (Opn7b: quantified areas: n = 46, parapinopsin: quantified areas: n = 68). **I)** Representative images of the injected BNST region showing cFos expression and parapinopsin or Opn7b distribution. ov: oval nucleus of the BNST, ad: anterodorsal nucleus of the BNST. **J)** Representative images of the VTA region (tyrosine-hydroxylase (TH) used as a marker for dopaminergic neurons) showing cFos expression in mice expressing either parapinopsin or Opn7b in the BNST. fr: fasciculus retroflexus. Scale bar = 300 µm Data are shown as means ± SEM. *p ≤ .050, **p ≤ .010, ***p ≤ .001 (unpaired Student’s t-test).

Since the BNST is interconnected with the VTA and both structures contribute to avoidance behavior, we next injected male mice with AAV expressing Opn7b or parapinopsin in the VTA and performed the same experiments as described above (Fig. 2A-F). Under low 2MT concentrations we found a decrease in the total distance moved (Fig. 2C), and an increase in freezing (Fig. 2F) in OPN7 injected mice compared to parapinospin injected mice. In addition, under high 2MT concentrations we observed a decrease in the total distance moved (Fig. 2C), a decrease in the time spent in the odor zone (Fig. 2E), and an increase in freezing in comparison to parapinopsin expressing mice (Fig. 2F). In addition, we observed an increase in time spent in the odor zone (Fig. 2E) if only water was presented in Opn7b in comparison to parapinopsin expressing mice. The experiments suggest that expression of Opn7b in the VTA increases avoidance related behavior in mice. We next quantified cFos-expressing neurons and found that Opn7b expression in comparison to parapinopsin in the VTA reduced their number by 61%. (Fig. 2G,I) and by 17% in the BNST (Fig. 2H,J).

**Figure 2:**
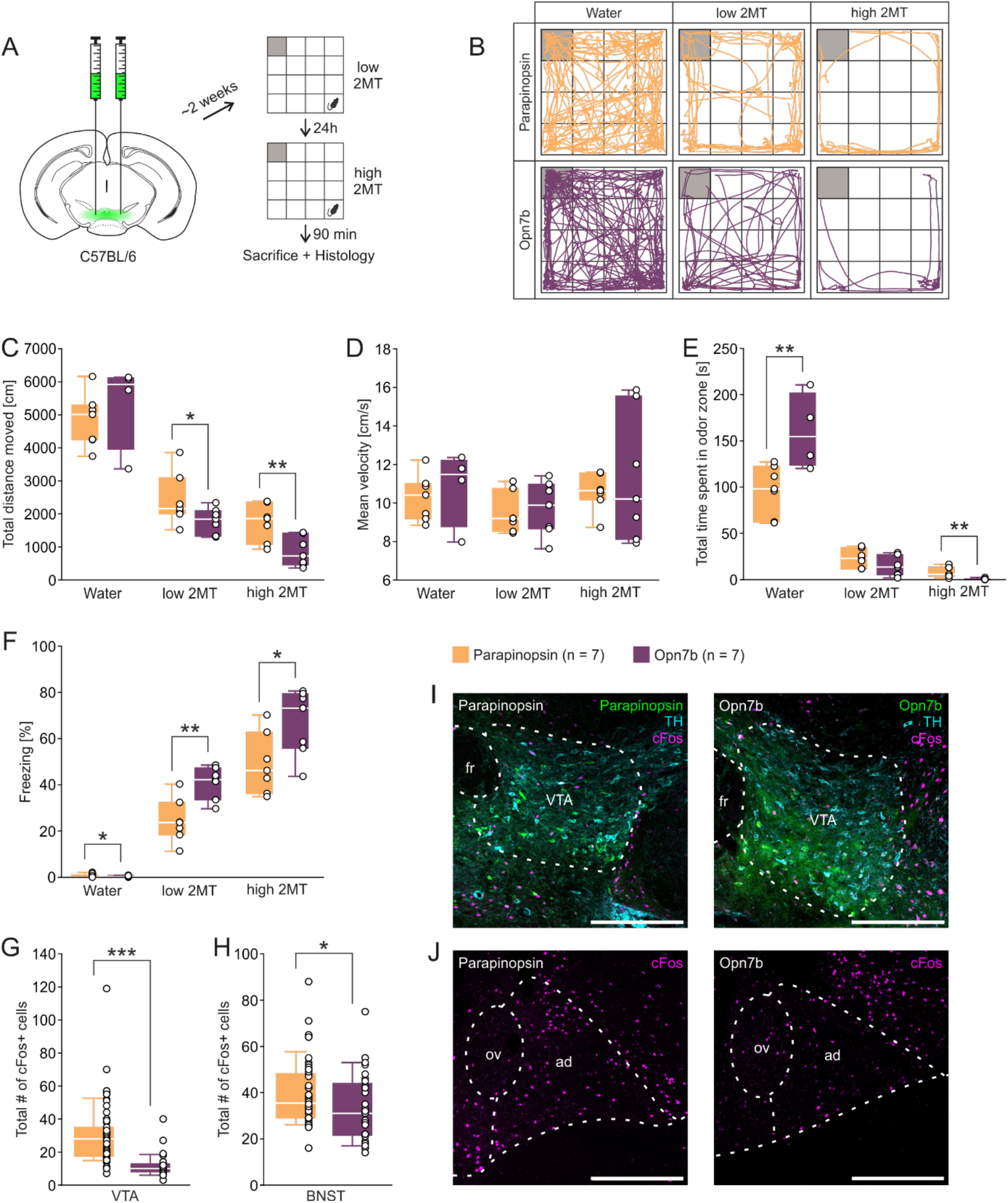
Injection of the constitutively active optogenetic tool Opn7b into the VTA reduces cFos expression in the VTA and BNSTad. **A)** Stereotactic injection of the viral constructs encoding either parapinopsin or Opn7b into the ventral tegmental area (VTA) of C57BL/6 mice followed by behavioral testing in the 2-methyl-thiazoline (2MT) paradigm with the grey square representing the odor zone. **B)** Exemplary trajectory plots of individual mice of both groups exposed to Water as well as low and high 2MT condition in an open field arena. **C)** Mice expressing Opn7b in the VTA show a decreased total distance moved compared to mice expressing parapinopsin in the low 2MT condition (t_(12)_ = 1.90, p = .041) as well as the high 2MT condition (t_(12)_ = 3.20, p = .004). **D)** No differences between groups in the mean velocity in all conditions were found. **E)** Opn7b injection increased the time spent in the odor zone during the Water condition (t_(9)_ = -3.13, p = .006) and decreased it during high 2MT exposure (U = 1.00; z = -3.071, p = .001, r = 0.82). **F)** Mice expressing Opn7b in the VTA showed less freezing in the Water condition (t_(9)_ = 2.15, p = .030) but greater freezing percentages when exposed to low (t_(12)_ = -3.72, p = .001) and high 2MT compared to mice expressing parapinopsin (t_(12)_ = -2.41, p = .016). **G)** The mice expressing Opn7b (quantified areas: n = 33) show a decrease in the total number of cFos immunoreactive cells in the VTA (U = 168.00; z = -6.472, p < .001, r = 0.68). (parapinopsin: quantified areas: n = 57). **H)** Mice expressing Opn7b (quantified areas: n = 29) in the VTA show a decrease in the total number of cFos immunoreactive cells in the anterodorsal nucleus of the bed nucleus of the stria terminalis (BNSTad) (U = 470.00; z = -2.145, p = .032, r = 0.25) compared to mice expressing parapinopsin (quantified areas: n = 46) in the VTA. **I)** Representative images of the injected VTA region (tyrosine-hydroxylase (TH) used as a marker for dopaminergic neurons) showing cFos expression and parapinopsin or Opn7b distribution. fr: fasciculus retroflexus. Scale bar = 300 µm. **J)** Representative images of the BNST region showing cFos expression in control mice and mice expressing parapinopsin or Opn7b in the VTA. ov: oval nucleus of the BNST, ad: anterodorsal nucleus of the BNST. Scale bar = 300 µm. Data are shown as means ± SEM. *p ≤ .050, **p ≤ .010, ***p ≤ .001 (unpaired Student’s t-test).

## Discussion

The main goal of this study was to determine whether the constitutive activation of the Gi/o pathway by Opn7b suppresses cFos expression in neuronal circuits and if this suppression correlates with behavioral changes. Indeed, in comparison to the non-constitutively active parapinospin expression of Opn7b in the BNST and the VTA reduced cFos expression and increased odor-induced avoidance and fear behavior. The reduction in cFos expression can be explained by the constitutive activation of the G_i/o_ pathway mediated by Opn7b (Karapinar et al., 2021). Constitutive activity of Opn7b stabilizes the resting membrane potential, while deactivation of Opn7b-mediated constitutive activity synchronizes neuronal network activity (Karapinar et al., 2021). The activation of the G_i/o_ pathways seems to be important to inhibit the activation of the immediate early genes in neurons. A reduction in cFos activity has also been observed when G_i/o_ coupled inhibitory DREADDs have been expressed and activated in the central nervous system (Bruzsik et al., 2021; Tiwari et al., 2022).

Opn7b-mediated reduction of cFos expression in the BNST and VTA increased avoidance and fear-related behaviors, compared to parapinopsin, in response to potential threats. Both brain areas are involved in encoding avoidance behavior and are reciprocally connected. For example, VTA dopaminergic neurons project into the dorsomedial nucleus of the BNST (Lebow and Chen, 2016), while peptidergic and GABAergic neurons innervate in particular dopaminergic neurons in the VTA (Soden et al., 2022). Thus, it is likely that Opn7b-mediated reduced constitutive baseline activity of GABAergic and dopaminergic projections from the BNST and VTA drives the modulation of innate fear responses. In fact, chemogenetic inhibition of GABAergic neurons in the BNST using G_i/o_-coupled DREADDs increased avoidance behavior and freezing in the presence of cat odors and 2MT (Bruzsik et al., 2021). On the other hand, dopaminergic activity and release in the VTA is increased by cued warning signals leading to avoidance behavior (Duvarci, 2024). We observed, that Opn7b expression in the VTA leads to 61% decrease in cFos positive neurons and a 17% reduction in the BNST. Therefore, constitutively reducing the activity of VTA-dopamine release may silence specific GABAergic populations to increase innate avoidance behavior by disinhibiting stress-promoting circuits. Another possibility is that Opn7b increases the signal-to-noise ratio in the BNST and VTA for incoming threat stimuli leading to strong DA release due to reduced baseline activity of Opn7b-expression cells.

An important question is whether the decreased cFos expression can directly correlate with changes in GPCR-signaling and network activity and avoidance behavior. In our experiments the mean cFos expression in the VTA is negatively correlated with time to freezing (supplemental Fig. 1), which is consistent with the observations that higher DA activity in the medial VTA is negatively correlated with freezing in extinction learning (Cai et al., 2020). A few studies investigated if the inhibition of cFos expression directly alters behavior. For example, inhibiting cFos expression through antisense oligonucleotides in rats resulted in deficits in spatial long-term memory (Gallo et al., 2018). Additionally, various drugs, such as diazepam, or chemogenetic inhibition decreased cFos expression in areas associated with stress, anxiety and avoidance behavior including the BNST (De Medeiros et al., 2005). These experiments suggest that cFos expression can be linked to specific behavioral responses strengthening the possibility of utilizing Opn7b-mediated suppression of cFos for investigating pathway-specific regulation of behavior.

In conclusion, the non-visual opsin Opn7b can be used to increase constitutive G_i/o_ activity in specific brain regions and specific cell-types without supplying exogenous ligands or drugs and is therefore a tool to investigate the role of constitutive GPCR activity in network function and disease.

## Author contribution

Conceptualization, I.S., S.H., M.H., and K.S.; Methodology and Investigation, H.B., H.S., M.W., J.B., and B.M.; Data analysis, H.B., H.S., M.W., and J.B.; Funding acquisition, I.S., S.H., M.H., and K.S.; Supervision, I.S., S.H., M.H., and K.S.; Writing, H.B., H.S., M.W., I.S., S.H., M.H., and K.S.

## Funding

Open Access funding enabled and organized by Projekt DEAL.

### Acknowledgments

We thank Winfried Junke, Stephanie Krämer, Manuela Schmidt, Gina Hillgruber, Elli Buschtöns, Petra Knipschild, Margareta Möllmann, Stefan Dobers, and Henning Knoop and the RUB mechanical shop for technical support. This work was supported by Deutsche Forschungsgemeinschaft (DFG) grants: Project ID 316803389 - SFB 1280, K.S., S.H., M.H. (Subproject A07, A18); Project number 492434978 - GRK 2862/1, Sub-projects (01, S.H.; 07, K.S.; 09, I.S.).

---

**Supplementary figure 1:**
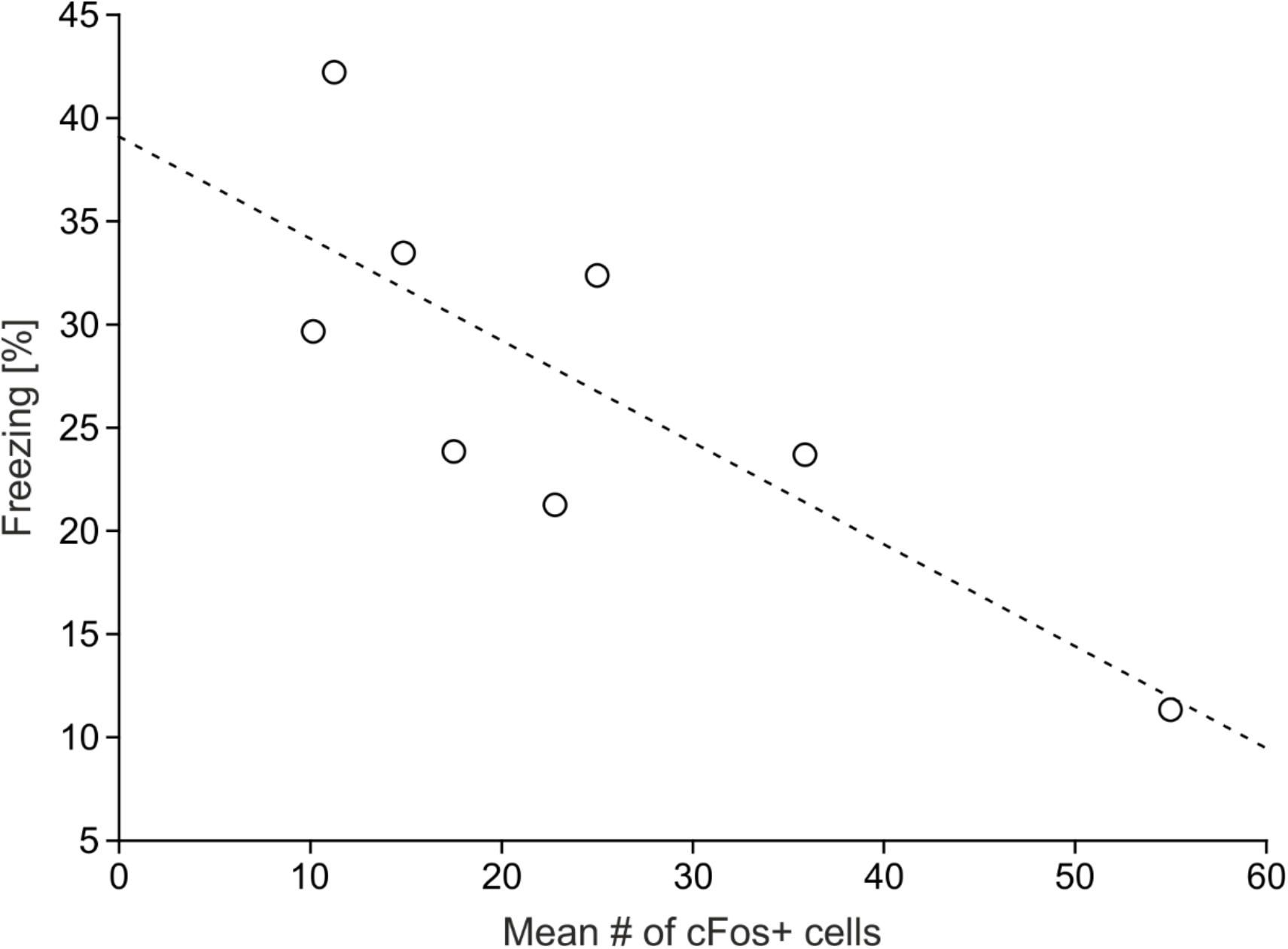
Negative correlation between freezing behavior and VTA neuronal activity during the low 2MT condition. Scatter plot showing a significant negative correlation (R = -80, p = .0017) between the percentage of freezing during the low 2MT condition and the total number of cFos-positive cells in the ventral tegmental area (VTA). Each data point represents an individual subject. The line represents the linear regression fit.

**Supplementary table 1:**
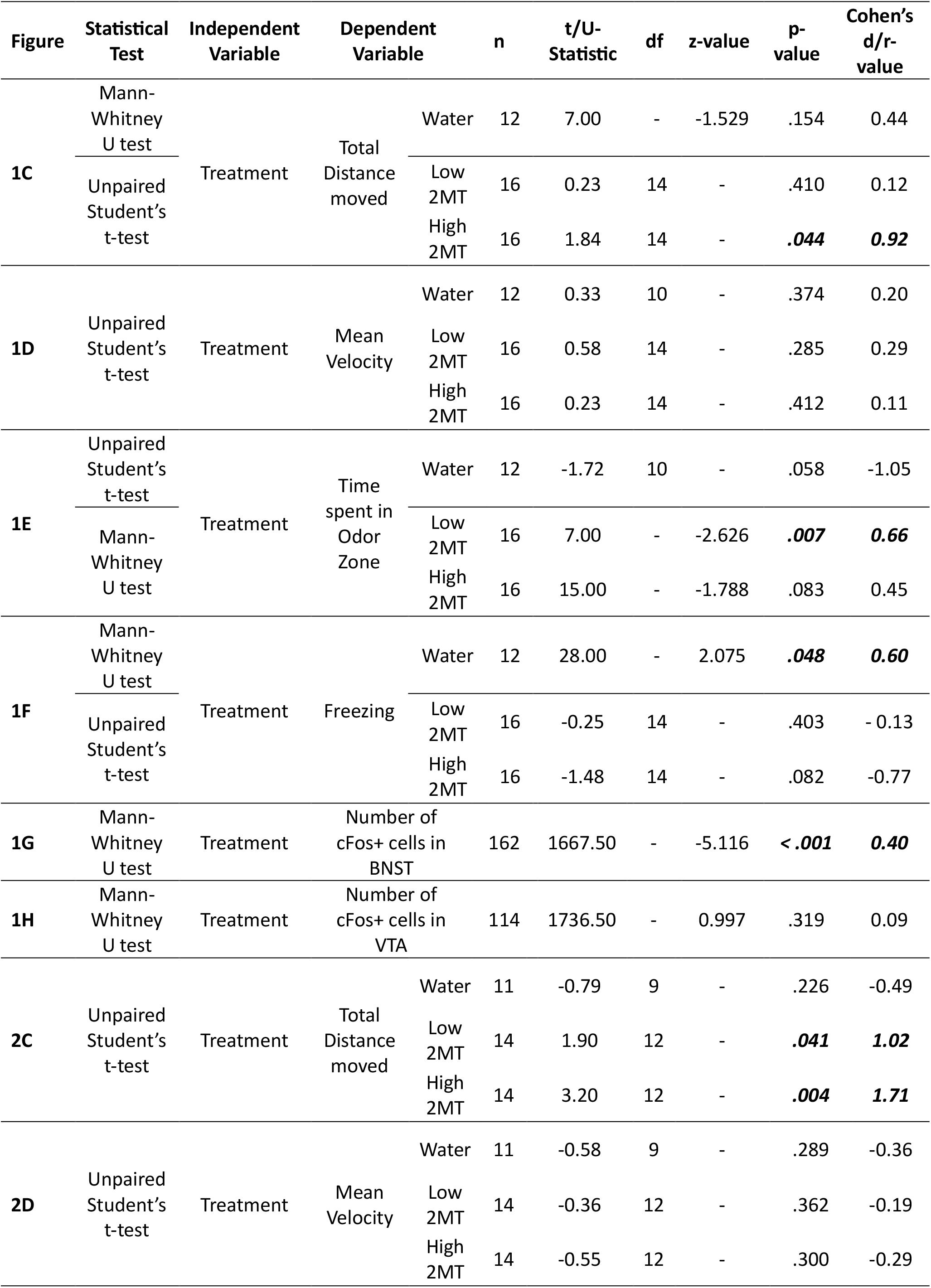
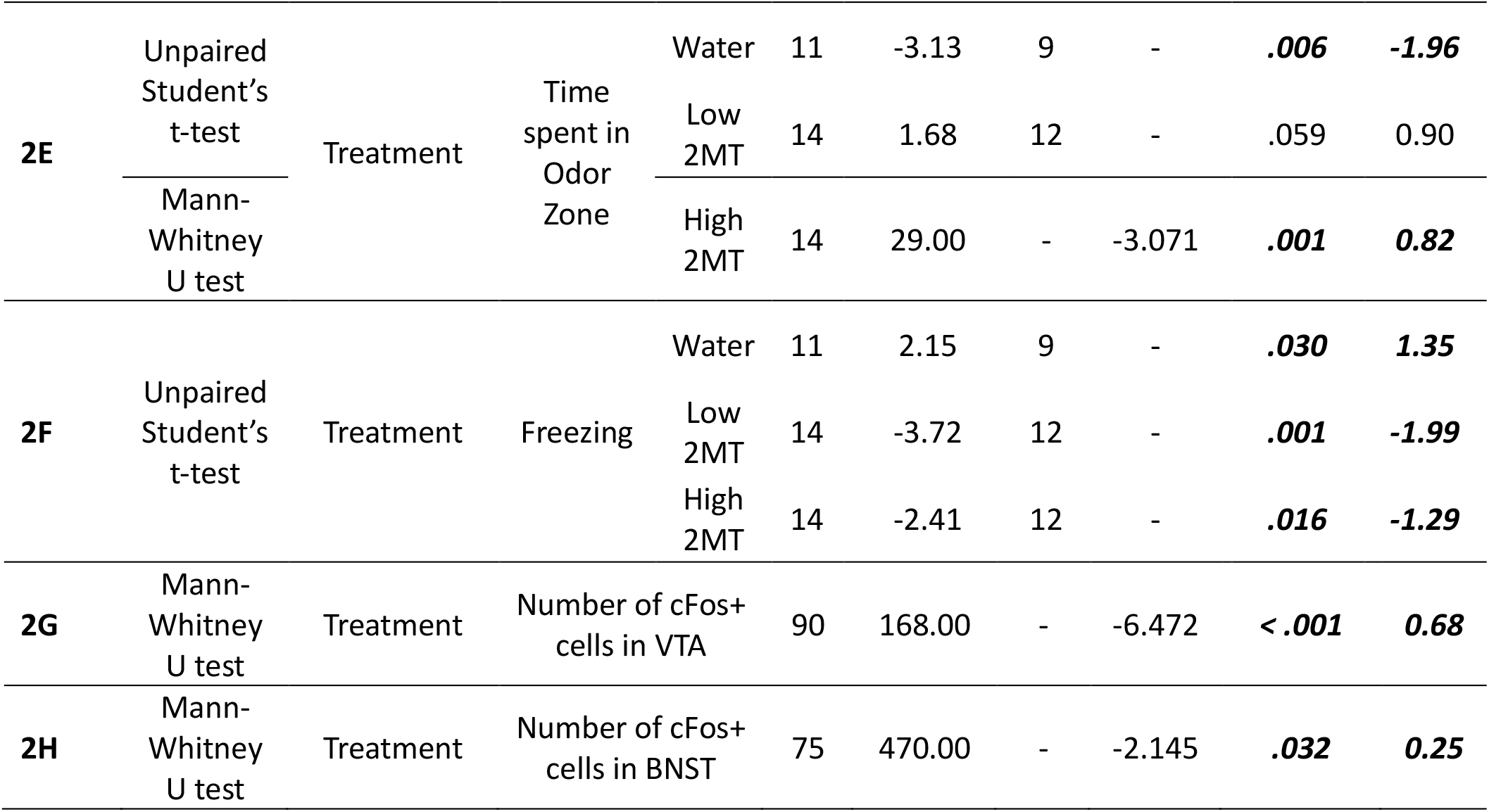
Statistical procedures and results. Statistical comparison of parapinopsin or Opn7b injection (treatment) in the BNSTad **(1C-1H)** and VTA **(2C-2H)** regarding behavioural parameters and cFos expression.

